# SNR-efficient distortion-free diffusion relaxometry imaging using ACcelerated Echo-train shifted EPTI (ACE-EPTI)

**DOI:** 10.1101/2021.08.27.457992

**Authors:** Zijing Dong, Fuyixue Wang, Lawrence Wald, Kawin Setsompop

## Abstract

**Purpose:** To develop an efficient acquisition technique for distortion-free diffusion MRI and diffusion-relaxometry.

**Methods:** A new ACcelerated Echo-train shifted Echo-Planar Time-resolved Imaging (ACE-EPTI) technique is developed to achieve high-SNR, distortion- and blurring-free diffusion and diffusion-relaxometry imaging. ACE-EPTI employs a newly designed variable density spatiotemporal encoding with self-navigation capability, that allows submillimeter in-plane resolution using only 3-shot. Moreover, an echo-train-shifted acquisition is developed to achieve minimal TE, together with an SNR-optimal readout length, leading to ~30% improvement in SNR efficiency over single-shot EPI. To recover the highly accelerated data with high image quality, a tailored subspace image reconstruction framework is developed, that corrects for odd/even-echo phase difference, shot-to-shot phase variation, and the *B*_0_ field changes due to field drift and eddy currents across different dynamics. After the phase-corrected subspace reconstruction, artifacts-free high-SNR diffusion images at multiple TEs are obtained with varying T_2_* weighting.

**Results:** Simulation, phantom and *in-vivo* experiments were performed, which validated the 3-shot spatiotemporal encoding provides accurate reconstruction at submillimeter resolution. The use of echo-train shifting and optimized readout length improves the SNR-efficiency by 27-36% over single-shot EPI. The reconstructed multi-TE diffusion images were demonstrated to be free from distortion (susceptibility and eddy currents) and phase/field variation induced artifacts. These improvements of ACE-EPTI enable improved diffusion tensor imaging and rich multi-TE information for diffusion-relaxometry analysis.

**Conclusion:** ACE-EPTI was demonstrated to be an efficient and powerful technique for high-resolution diffusion imaging and diffusion-relaxometry, which provides high SNR, distortion- and blurring-free, and time-resolved multi-echo images by a fast 3-shot acquisition.

## Introduction

Diffusion MRI (dMRI) is a critical imaging tool that can probe tissue microstructure non-invasively by measuring the diffusion process of water molecules (1). The application of dMRI in clinical research and large-scale studies, such as human connectome project (HCP) (2,3) and UK biobank (4), has facilitated a better understanding of neurological diseases and brain connectivity. For decades, single-shot echo-planar imaging (ss-EPI) is the most commonly used acquisition method in dMRI, which is known for its fast acquisition speed and insensitivity to motion. However, ss-EPI suffers from image distortion and blurring due to *B*_0_-inhomogenity-induced phase accumulation and signal decay across different phase encoding (PE) signals, which becomes more prominent at higher spatial resolution, limiting the structural integrity and resolution. Therefore, conventional ss-EPI acquisition cannot fulfill the increasing need of dMRI to pursue higher spatial resolution to investigate more detailed brain structures and connectivity, such as at the gray-white matter interface or within the cortex (5,6). Moreover, the formation of a single image from a long readout in ss-EPI results in limited ability to acquire multiple echoes and in flexibility of TE selection, especially at high resolution. Such multi-TE information is highly desired for the emerging diffusion-relaxometry imaging to disentangle complicated tissue microstructures (7–11).

To reduce the distortion and blurring, parallel imaging (12–14) has been applied to ss-EPI, which can reduce the effective echo spacing (ESP) by undersampling along PE (15). The reduction of distortion/blurring is determined by the undersampling factor, which is around 2~4×, and a shorter echo-train length (ETL) can also be obtained. Since a high undersampling rate will lead to potential aliasing artifacts and large g-factor penalty with noise amplification, further reduction is limited using parallel imaging alone. As an alternative, multi-shot EPI (ms-EPI) methods have been developed (16–21). By acquiring different segments of the *K*-space in multiple shots, the effective ESP and ETL can be reduced, leading to effective distortion and blurring reduction without elevated g-factor. However, the level of distortion/blurring in ms-EPI is related to the number of shots, and non-negligible distortions still remain with practical protocols (e.g., 4-10 shots), especially for regions with strong susceptibility change such as the temporal lobe and the frontal lobe. In addition, the shot-to-shot phase variation caused by physiological movements during the application of diffusion gradients can lead to severe image artifacts, so navigator is required for the physiological motion correct (16,18,22), which may need extra scan time and compromise its acquisition efficiency over ss-EPI. To achieve distortion- and blurring-free dMRI, point-spread-function (PSF) encoding approach was developed and combined with ms-EPI (23–26). It acquires an additional encoding dimension in a multi-shot manner to measure the PSF along the PE direction, which is free from both distortion and blurring. Our recently work on tilted-CAIPI encoding pattern (27), was able to reduce the acquisition shots of PSF-EPI from more than 200 (full sampling) to only 7-9 for imaging at ~1 mm in-plane resolution, allowing high-quality distortion-free dMRI within practical scan time.

Echo-planar time-resolving imaging (EPTI) (28) is a recently developed novel ms-EPI method, that not only achieves distortion- and blurring-free imaging, but also provides time-resolved multi-echo multi-contrast images spaced at an ESP (~1 ms) apart to track the signal evolution for accurate relaxometry estimation. These markable features of EPTI rely on the use of a new time-resolving approach for continuous EPI readout, where the full *k-t* (frequency-echo) space is recovered from a highly-undersampled data acquired with an optimized spatiotemporal encoding scheme. The high acquisition efficiency and improved image quality for multi-contrast imaging of EPTI have enabled ultra-fast quantitative relaxometry (29,30) and the potential for diffusion-relaxometry imaging (31,32). However, in order to achieve high in-plane resolution (e.g., submillimeter) in 2D EPTI, 5-10 shots are still required for each image slice. For dMRI, the long acquisition time leads to higher sensitivity to inter-shot motion and less flexibility of diffusion encoding selection.

In this work, we present an efficient acquisition method for dMRI, named ACcelerated Echo-train shifted EPTI (ACE-EPTI), which achieves high-SNR, distortion- and blurring-free diffusion and diffusion-relaxometry imaging at submillimeter resolution in just 3 shots. ACE-EPTI employs a newly designed variable density spatiotemporal encoding with self-navigation capability, to better take the advantage of the locally low-rank (LLR) constraint in addition to the spatiotemporal correlation in the reconstruction. Moreover, enabled by the flexibility of time-resolved imaging, an echo-train-shifted acquisition is developed to minimize the TE, together with the SNR-optimized readout length, leading to 27-36% improvement in SNR efficiency over the conventional ss-EPI acquisition in the examined protocols. To recover the highly undersampled *k-t* data, a tailored subspace image reconstruction framework is developed for ACE-EPTI, which corrects for odd/even echo phase difference, shot-to-shot phase variation using self-navigators, as well as the *B*_0_ field changes across different dynamics/directions. After the phase-corrected LLR subspace reconstruction, artifacts-free high-SNR diffusion weighted images (DWIs) at multiple TEs with varying T_2_* weighting can be obtained from a single ACE-EPTI dataset acquired in 3 shots. Simulation, phantom and *in-vivo* experiments are performed to validate: i) accurate reconstruction at submillimeter resolution using the 3-shot spatiotemporal encoding, ii) higher SNR efficiency using ACE-EPTI with echo-train shifting and optimized readout length than ss-EPI at different resolutions, iii) anatomical integrity without both susceptibility and eddy-current induced distortion, iv) artifact-free multi-TE DWIs recovered by the proposed phase-corrected subspace reconstruction framework. These improvements of SNR and image quality using ACE-EPTI are also demonstrated to provide improved diffusion tensor imaging (DTI) and efficient acquisition for diffusion-relaxometry analysis through *in-vivo* dMRI experiments.

## Theory

### Echo-train shifted acquisition with flexible readout length

The time-resolving approach in EPTI allows image recovery at each echo time point across the EPI readout instead of a single image from combining data across the full echo train (28). This provides the flexibility of the readout to start at any TE and is still able to recover all the desired contrast-weighted images as long as they are sampled within the readout window. In order to minimize T_2_ decay and achieve the highest possible SNR, the readout echo-train is shifted to start right at the spin-echo (SE) time point after the diffusion gradients in the ACE-EPTI sequence as shown in Fig. 1a, achieving minimal TE similar to a half partial-Fourier acquisition but without any loss of spatial resolution. A series of T_2_*-weighted images can be resolved following the T_2_-weighted at the SE time-point, with a time increment of an ESP. These multi-echo images can be either used separately for diffusion-relaxometry, or combined to a single image with higher SNR, where the strong signal dephasing area can be well compensated using a simple sum-of-squares (SoS) combination to increase the weights of the early echoes. The reduction in TE achieved with ACE-EPTI over ss-EPI would increase with increasing spatial resolution, making it an efficient acquisition for high-resolution dMRI.

**Figure 1:**
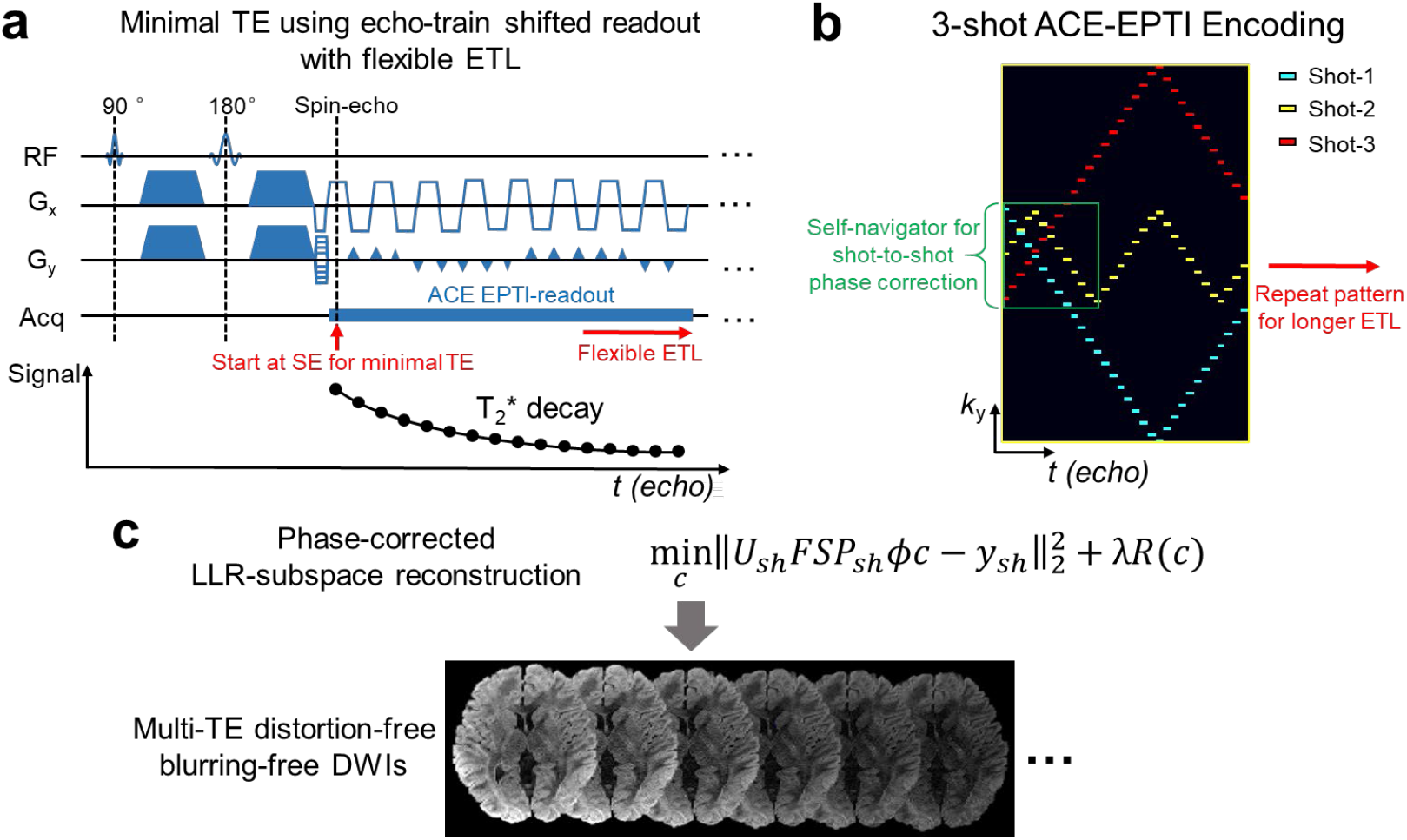
Illustration of ACE-EPTI acquisition, encoding, and reconstruction. (a) The sequence diagram of ACE-EPTI acquisition which utilizes echo-train shifting to minimize TE. The continuous readout starts at the spin-echo point and can be played with flexible echo-train length (ETL) to track T_2_* decay. (b) The optimized variable density 3-shot spatiotemporal encoding pattern, with a portion of low-frequency signals of each shot used for self-navigated shot-to-shot phase variation correction (green window). (c) Phase-corrected LLR-subspace reconstruction to resolve the multi-echo distortion- and blurring-free images from the highly-undersampled *k-t* data.

In addition to minimizing the TE, the time-resolving approach also enables flexible ETL selection since the image resolution is determined by the design of *k-t* encoding instead of the ETL, and a longer ETL will not cause distortion or blurring but provides more echoes and signals. This allows us to optimize the ETL to further improve the SNR efficiency, by acquiring more echoes for better noise averaging (33), or reducing the ETL to avoid acquiring low signal echoes and to reduce the TR.

### 3-shot self-navigated k-t encoding

The optimized 3-shot spatiotemporal encoding is presented in Fig. 1b, which provides further acceleration in the *k-t* space to reduce the acquisition time of each volume for improved motion robustness and the flexibility of more diffusion encodings. The fast 3-shot encoding in ACE-EPTI is designed with: (i) complementary *k*_y_ (PE) sampling along *t* to control aliasing, (ii) denser sampling at central k-space and early TE to improve SNR and allow a more accurate locally low-rank constrained reconstruction, (iii) small *k*_y_ changes between neighboring time points to limit the gradient blip size, to reduce the ESP and potential eddy currents, (iv) low-frequency signals acquired in each shot (green window) for self-navigated shot-to-shot phase variation estimation to correct for physiological motion. These optimizations of the spatiotemporal encoding allow accurate reconstruction of time-resolved multi-contrast images from the highly-accelerated *k-t* data by exploiting the strong temporal signal correlation, multi-channel receiver coil and LLR property in the spatial/coefficients domain. Note that the pattern shown in Fig. 1b is a low resolution example, with a *k*_y_ matrix size of 128 and a maximum change of 4 *k*_y_ between neighboring time points. The encoding pattern will be scaled up accordingly for higher spatial resolution cases, with more echoes acquired in each up and down sections, proportional to the increased matrix size, and with the pattern repeated for longer ETL.

For larger volumetric coverage, the designed 3-shot encoding for ACE-EPTI is also combined with blipped-CAIPI encoding (34) along *k*z for simultaneous multi-slice (SMS) acquisition of the ACE-EPTI sequence. The same standard blipped-CAIPI encoding was applied for all the 3 shots, and multi-slice images can be recovered with multiple TEs by the subspace reconstruction described below.

### Time-resolved subspace reconstruction

A phase-corrected LLR subspace reconstruction (Fig. 1c) is used to resolve the highly undersampled *k-t* data to multi-contrast images. The main idea in subspace reconstruction is that if the temporal dimension of an image series is low rank, all the possible signal evolutions can be represented accurately by a vector subspace (35–38). The basis vectors (*ϕ*) for the subspace can be obtained from the simulated signal evolutions based on the signal model. By representing the signal evolutions with the subspace model, only the coefficients (*c*) of the subspace bases need to be estimated in each voxel to recover the full time series, instead of estimating all the time points independently, leading to significantly reduced number of unknowns and improved conditioning in the reconstruction. In addition to utilizing the temporal correlation through subspace modeling, additional constraints such as LLR can also be applied to further improve the accuracy and SNR by exploiting the low-rankness in spatial/coefficients domain (36,37,39). The subspace approach has been used in previous EPTI works and has proven to provide superior performance for highly accelerated data (29,30,37). For ACE-EPTI, the subspace reconstruction is modified to account for the shot-to-shot phase variation in dMRI acquisition:

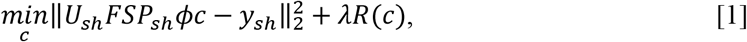

where *c* is the coefficient maps of the subspace bases *ϕ*, *P_sh_* is the phase evolution across the EPI readout of the *sh*-th shot including the background and *B*_0_-inhomogneity-induced phase, *S* is the coil sensitivity map, *F* represents the Fourier transform operator, *U_sh_* is the *k-t* undersampling mask corresponding to the acquired *sh*-th shot signals, *y_Sh_*. *R*(*c*) denotes the LLR regularization applied on *c* with a control parameter *λ*. The subspace bases are generated from simulated signals based on the Bloch equation, and all the other parameters including coil sensitivity (*S*) and phase maps (*P_Sh_*) can be calculated from a short *k-t* calibration scan and phase estimation process that will be illustrated below. After solving the only unknown *c* in Eq. [1], the multi-contrast image series can be recovered as *ϕc*, free of distortion and blurring artifacts.

### Odd/even echo phase correction, shot-to-shot phase correction, and B_0_ field updating

Accurate phase modeling of the EPTI signals is critical to ensure the reconstruction accuracy using the subspace approach. To achieve this, a phase-correction framework is developed as shown in Fig. 2 to provide accurate phase modeling of the ACE-EPTI data, including odd/even echo phase correction, shot-to-shot phase variation correction, and dynamic *B*_0_ field updating.

**Figure 2:**
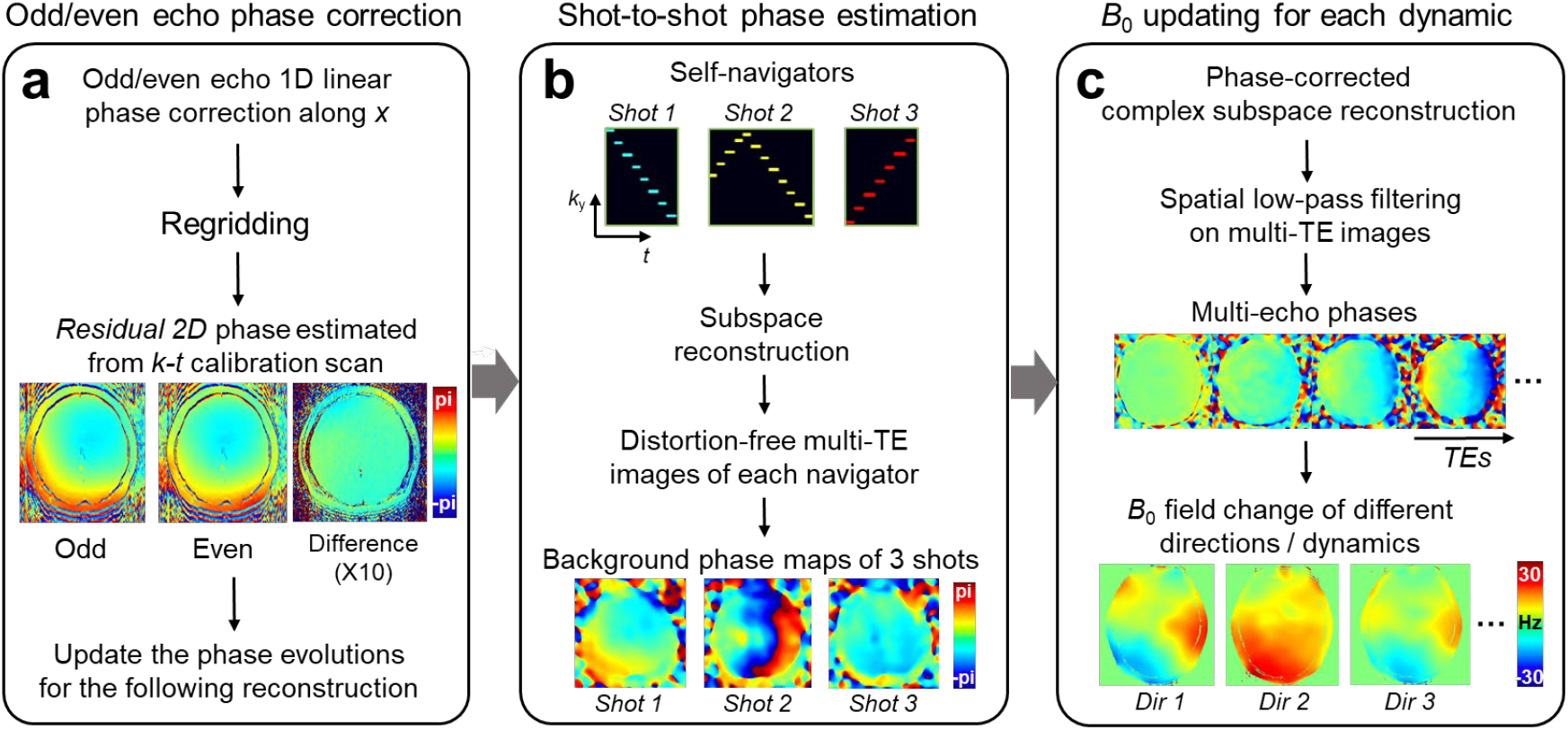
Illustration of the phase correction framework. It includes correction for (a) odd/even echo phase difference, (b) shot-to-shot phase variation, and (c) dynamic *B*_0_ changes of different volumes. (a) The phase difference between odd and even echoes is estimated from the *k-t* calibration scan. The 1D linear phase difference along *x* is corrected first before regridding, and the residual 2D phase difference is then added into the phase term in the subsequent subspace reconstruction. (b) The low-frequency signals from the 3-shots ACE-EPTI serve as self-navigators and are reconstructed to obtain distortion-free images by subspace reconstruction. The shot-to-shot phase variation is then estimated from the background phase at the SE, and modeled in the phase-corrected subspace reconstruction. (c) A subspace reconstruction is performed with complex bases to estimate the *B*_0_ map for each dynamic before the final image reconstruction. A low-pass filter is applied on the reconstructed multi-echo images to improve the SNR assuming that the *B*_0_ change should be smooth, and the *B*_0_ changes are estimated by a temporal linear fitting of the multi-echo phases.

The phase difference between odd and even echoes caused by eddy currents and gradient delay is well-recognized in EPI readout, which can result in ghosting artifacts in conventional EPI images. To mitigate this issue, 1D navigators are typically acquired after each excitation pulse to measure the 1D phase difference along the readout (*x*) direction between the odd and even echoes, in which most of the eddy currents appear. Although the designed ACE-EPTI sequence avoids the use of such navigator acquisition to achieve minimal TE, the odd/even echo phase difference can be well estimated from the *k-t* calibration scan. Specifically, the 2D phase differences between odd and even echoes of each slice are estimated using the fully sampled *k-t* calibration by extract the background phase of odd and even echoes respectively after removing the fitted *B*_0_-induced phase. As shown in Fig. 2a, the 1D linear phase difference along *x* is first corrected in the raw signals before regridding (ramp sampling correction), and after regridding, the residual 2D phase differences are corrected in the phase evolution term P using the different background phases of odd and even echoes.

The shot-to-shot phase variation due to physiological motion is estimated using the self-navigators in ACE-EPTI as shown in Fig. 2b. A subspace reconstruction is applied for the navigator of each shot separately, to recover low-resolution images without distortion. Background phase maps of the 3 shots are extracted from the reconstructed images at the SE point, corresponding to the shot-to-shot phase variations. To correct for the phase variations in the following subspace reconstruction, the estimated background phase of each shot is added into the phase term *P*_sh_ in Eq. [1].

An initial *B*_0_ map is estimated from the *k-t* calibration scan, however, the field inhomogeneity can change in the dMRI acquisition due to field drift or eddy currents. To provide accurate *B*_0_ map for each dynamic/volume (every 3 shots), the *B*_0_ map is updated using an initial subspace reconstruction before the final image reconstruction (Fig. 2c). In this initial reconstruction, complex signal bases are generated that can represent the signal evolution as well as additional phase evolutions from a temporal *B*_0_ change ranging from −50 to 50 Hz. A larger number of bases is required to estimate this addition phase evolution, so a lower SNR in this reconstruction is expected when compared to that of the final subspace reconstruction which will utilize a reduced basis set that do not account for this additional phase evolution. By applying a spatial low-pass filter to the reconstructed multi-echo images, high SNR phase maps can still be obtained to fit for an accurate *B*_0_ change estimate that can be used in the final subspace reconstruction. The updated *B*_0_ map is also used in the pre-reconstruction of the next dynamic data to ensure the difference between the initial *B*_0_ map and data is within ±50 Hz.

## Methods

All of the experiments were performed on a Siemens Prisma 3T scanner with a 32-channel head receiver coil (Siemens Healthineers, Erlangen, Germany). In-vivo data were acquired on healthy volunteers (N=2) with a consented institutionally approved protocol.

### Evaluation of the 3-shot encoding by retrospective undersampling test

Fully-sampled *k-t* EPTI data were acquired and retrospectively undersampled with different spatiotemporal encoding patterns to demonstrate the reconstruction accuracy that can be achieved using the designed 3-shot ACE-EPTI encoding. A spin-echo ACE-EPTI sequence without diffusion encoding was used to acquire this full *k-t* data at a spatial resolution of 0.86 × 0.86 × 3 mm^3^, and echo-shifting was used to achieve a starting TE of 31 ms. The matrix size was 256 × 255 × 16 × 64 (*k*_x_ × *k*_y_ × *slice* × *echo*), TE range was 31 – 93 ms, ESP = 0.97 ms, 255 shots were acquired with a TR of 3 s, and the total acquisition time was ~13 minutes. The fully-sampled data were then subsampled with 3 different patterns: 5-shot EPTI, 3-shot EPTI, and 3-shot ACE-EPTI. All data were reconstructed using LLR subspace reconstruction to evaluate their accuracy by comparing with the fully-sampled reference.

### Optimization of ETL using Monte-Carlo simulation

To take advantage of the flexible ETL in ACE-EPTI and optimize it for high SNR efficiency, a Monte-Carlo simulation was performed to evaluate the SNR performance under different ETLs. ACE-EPTI *k-t* data were simulated using pre-acquired quantitative maps, including proton density (PD), T_2_, T_2_*, and *B*_0_. A 32-channel dataset was generated using previously acquired coil sensitivity maps and undersampled by the 3-shot ACE-EPTI encoding pattern. Random noise (SNR = 20) was then added to the simulated *k-t* data. Data with 30 replicas were simulated and reconstructed to calculate the SNR based on the temporal SNR equation (mean magnitude / standard deviation of the dynamic images). The data were simulated at a moderate 1.5-mm in-plane resolution with an ESP of 0.66 ms in this evaluation, that will also be used later for *in-vivo* dMRI in additional to the high in-plane resolution protocol. The starting TE was set to 32 ms, close to the real ACE-EPTI acquisition. Image SNRs at various ETLs from 33 ms (50 echoes) to ~100 ms (150 echoes) were analyzed.

To determine the optimal ETL for highest efficiency, SNR efficiency was calculated based on the obtained SNRs as:

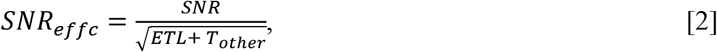

where *ETL* is the readout time of each slice, and *T*_other_ is the total event time for the other components in the sequence, including RF pulses, gradients (diffusion, pre-rewinder, spoiler, etc.,) and fat saturation, which was set to 65 ms. Note that the impact of the differences in TRs on T1 recovery due to differences in ETL was not considered in the SNR-efficiency calculation. This effect should be minimal for acquisitions with sufficiently long TRs.

### SNR analysis in phantom and in-vivo experiments

Both phantom and *in-vivo* data were acquired using the ACE-EPTI sequence to demonstrate its ability to provide improved SNR-efficiency when compared to conventional ss-EPI. Two protocols were tested at different resolutions: 1.5-mm isotropic and 0.86 × 0.86 × 3 mm^3^. SNR maps were calculated from the acquired non-DW 40-dynamic datasets for both ACE-EPTI and ss-EPI. The TEs used for both ACE-EPTI and ss-EPI were adjusted close to the corresponding dMRI acquisitions at *b* = 1000 s/mm^2^.

The key acquisition parameters for the 1.5-mm isotropic ACE-EPTI protocol were: matrix size = 160 × 159 × 28 × 85 (*k*_x_ × *k*_y_ × *slice* × *echo*), 3-shot acquisition, ESP = 0.66 ms, TE_range_ = 32 – 87 ms, TR = 3 s. The acquisition parameters for the 0.86-mm ACE-EPTI protocol was: matrix size = 256 × 255 × 12 × 64 (*k*_x_ × *k*_y_ × *slice* × *echo*), 3-shot acquisition, ESP = 0.97 ms, TE_range_ = 31 – 92 ms, TR = 3 s. The ss-EPI data were acquired at the same spatial resolutions and TR, where 3 averages with matching total acquisition time to that of ACE-EPTI were acquired to provide a fair comparison. The other parameters were: partial Fourier = 6/8, in-plane acceleration factor = 4, TE = 41 ms for the 1.5-mm protocol, and 57 ms for the 0.86-mm protocol.

A *k-t* calibration pre-scan was acquired before each of the ACE-EPTI scans, to estimate the *B*_0_ and coil sensitivity maps for ACE-EPTI reconstruction. The full *k-t* calibration data were acquired using a 2D GE EPTI sequence, with the same FOV and echo spacing as the imaging scan, but with a lower resolution, 35 × 150 × 7 (*k*_x_ × *k*_y_ × *echo*), and a much shorter TR (~25 ms per slice), resulting in a total acquisition time of ~ 1 minute.

### Diffusion acquisition at 0.86-mm in-plane resolution

*In-vivo* whole-brain dMRI data were acquired at 0.86-mm in-plane resolution using the 3-shot ACE-EPTI sequence to demonstrate the effectiveness of the proposed phase correction framework, the improved image quality with no distortion and artifacts, as well as reliable DTI analysis.

The diffusion ACE-EPTI data were acquired with the following parameters: 32 directions at *b* = 1000 s/mm^2^ together with 6 *b* = 0 s/mm^2^ volumes, in-plane resolution = 0.86 × 0.86 mm^2^, slice thickness = 3 mm, FOV = 220 × 220 × 126 mm^3^, matrix size = 256 × 255 × 42 × 64 (*k*_x_ × *k*_y_ × *slice* × *echo*), multi-band factor (MB) = 2, number of signal averages (NSA) = 3, ESP = 0.97 ms, TE_range_ = 38 – 99 ms, TR = 3.5 s, total acquisition time = ~20 minutes. Results before and after the proposed phase corrections were reconstructed to demonstrate the necessity of the odd/even echo phase correction, shot-to-shot phase variation correction and *B*_0_ updating in order to obtain artifacts-free high-quality DWIs. Moreover, T_2_* maps were fitted using data with and without diffusion weighting, together with TE-dependent Fractional anisotropy (FA) maps and orientation distribution functions (ODFs), to validate the potential use of multi-TE images provided by ACE-EPTI for diffusion-relaxometry.

Diffusion data were also acquired using ss-EPI with a matched scan time for comparison at the same spatial resolution. The same diffusion directions with NSA = 9 were acquired with the same TR (3.5 s with MB 2), while the TE was 64 ms with a partial Fourier factor of 6/8 and in-plane acceleration factor of 4. To obtain a distortion-free reference to evaluate the ACE-EPTI and ss-EPI images, TSE data were also acquired with the same spatial resolution.

### Multi-shell diffusion acquisition at 1.5-mm isotropic resolution

To further evaluate the ability of ACE-EPTI for diffusion-relaxometry imaging, in-vivo diffusion data at 1.5-mm isotropic resolution were acquired at *b* = 1000 s/mm^2^ and 2000 s/mm^2^. 3-shot ACE-EPTI acquisition was used with a MB factor of 3. The other imaging parameters were: 32 diffusion directions at *b* = 1000 s/mm^2^, and *b* = 2000 s/mm^2^, together with 12 *b* = 0 s/mm^2^ volumes (76 volumes in total), NSA = 2, FOV = 220 × 220 × 104 mm^3^, matrix size = 160 × 159 × 69 × 85 (*k*_x_ × *k*_y_ × *slice* × *echo*), ESP = 0.68 ms, TR = 4 s, TE_range_ = 54 – 111 ms, and TE was kept the same for all *b* values. The total acquisition time was ~30 minutes. The reconstructed multi-echo DWIs were then used to fit for T_2_* maps with different diffusion weightings, and the TE-dependent ODFs.

### Image reconstruction and post-processing

Subpace reconstruction was used for self-navigator (Fig. 2b), *B*_0_ updating (Fig. 2c), and the final image reconstruction (Fig. 1c). For navigator and image reconstruction, the subspace bases were generated by performing principal component analysis (PCA) of the simulated signals with different T_2_* decays (4 ms to 400 ms), and 3 bases were used resulting in an approximation error (mean nRMSE) < 0.2%. Additional *B*_0_ changes (−50 Hz to 50 Hz) were simulated to generate the complex bases used for the initial *B*_0_ updating subspace reconstruction, and 8 bases were used here to achieve the same approximation accuracy. A maximum number of iterations = 100 was set as the stop criterion, with a lambda of 0.01. The reconstruction algorithm was implemented based on BART toolbox (40,41) for computational acceleration.

Since ACE-EPTI is free from the eddy currents induced distortion, only rigid motions between images from different diffusion directions need to be corrected, which was done using the FLIRT (42,43) function in FSL (44,45). The skull was removed using BET (46) before image registration. FA maps were calculated using FSL, and orientation distribution functions (ODFs) were obtained and visualized in MRtrix3 (47). For ss-EPI data, POCS partial Fourier reconstruction was used and the in-plane undersampling was recovered using GRAPPA (14). For ss-EPI data, the ‘eddy’ function (48) was used to correct both the motion and eddy currents across different diffusion directions before calculating the FA maps.

All of the image reconstruction and post-processing were performed in MATLAB on a Linux workstation (CPU: Intel Xeon, 3.00GHz, 24 Cores; RAM: 512 GB; GPU: Quadro RTX 5000, 16 GB memory).

## Results

The reconstruction accuracy of different spatiotemporal encoding strategies is evaluated in a retrospective undersampling experiment as shown in Fig. 3a, including 5-shot EPTI, 3-shot EPTI, and 3-shot ACE-EPTI. Error maps and RMSEs were calculated by comparing the reconstruction images with the fully-sampled reference images after combining all the image echoes using SoS. The reconstructed image from the 3-shot ACE-EPTI shows markable improvement over the 3-shot EPTI, and achieves an accuracy close to the 5-shot EPTI acquisition. In addition, only noise-like error appears in the difference map of the 3-shot ACE-EPTI, without any noticeable artifacts.

**Figure 3:**
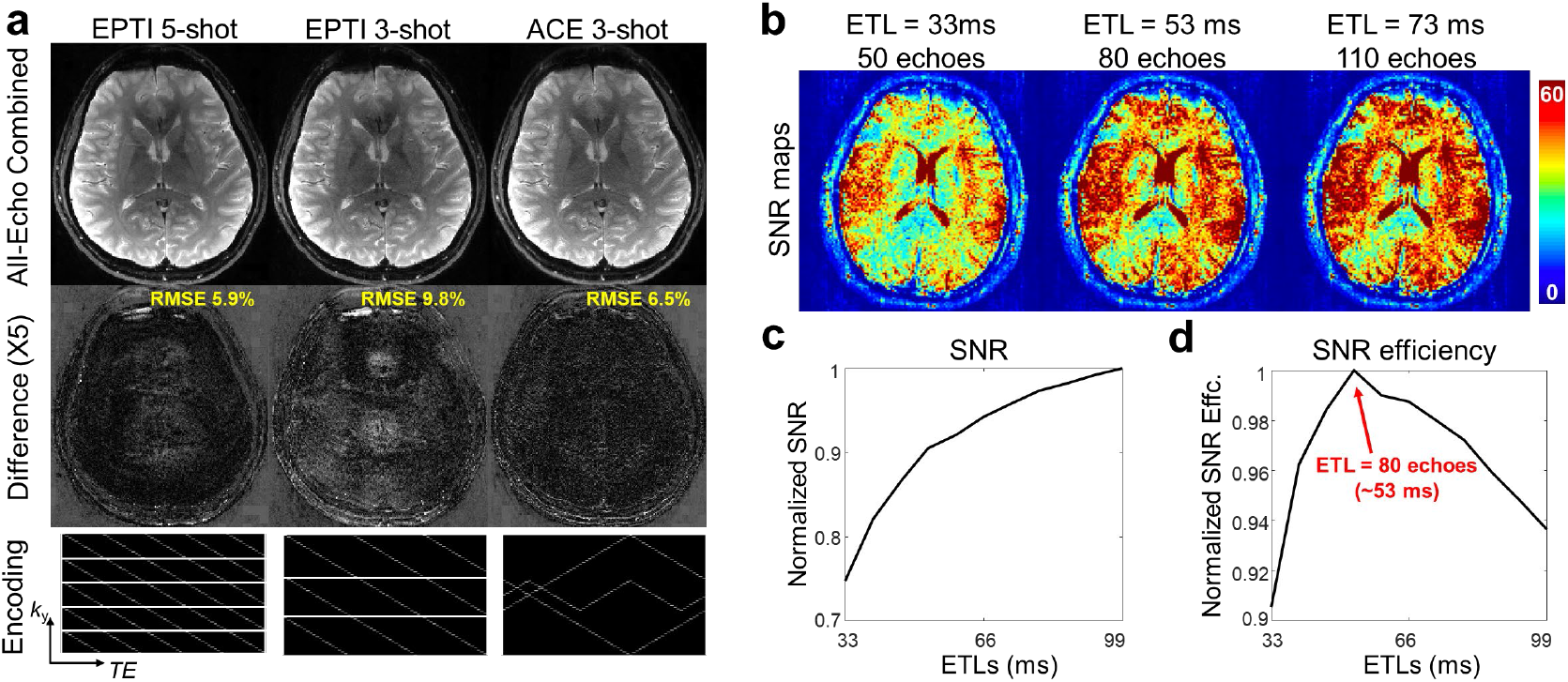
Evaluation of the optimized 3-shot ACE-EPTI encoding and optimization of the ETL. (a) Evaluation of the reconstruction accuracy of different spatiotemporal encoding schemes by retrospective undersampling, including 5-shot EPTI, 3-shot EPTI, and 3-shot ACE-EPTI. All-echo combined (using SoS) images are shown (top row), together with the error maps (middle row), and encoding patterns (bottom row). (b) Example SNR maps calculated from the Monte-Carlo simulation at ETL = 33 ms, 53 ms and 73 ms. (c) The normalized mean SNRs at different ETLs from 33 ms to 99 ms. (d) The normalized SNR efficiency. Highest SNR efficiency is achieved at an ETL of 53 ms (80 echoes, ESP = 0.66 ms) in the simulation.

Using the optimized 3-shot pattern, the SNR and SNR-efficiency of ACE-EPTI with different ETLs were analyzed in the Monte-Carlo simulation as shown in Fig. 3b-d. Figure 3b presents the SNR maps from 3 representative ETLs, calculated from the reconstructed images with 30 replicas at ETL = 33 ms, 53 ms and 73 ms. The normalized mean SNRs at different ETLs from 33 ms to 99 ms are plotted in Fig. 3c, showing that the SNR increases with longer ETL, but the rate of increase is slower at longer ETLs due to the reduced signals at the later TEs. Figure 3d plots the normalized SNR efficiency calculated based on Eq. [2]. An ETL of 53 ms (80 echoes with ESP = 0.66 ms) achieves the highest SNR efficiency in the analysis, and the result suggests that any ETL in the range of 45 to 70 ms can provide good SNR-efficiency, close to that of the optimal case. Since the impact of T1 recovery was not considered in the calculation, the SNR efficiency of a longer ETL can be underestimated if the TR is not very long. Hence, we chose a slightly longer ETL, around 56-62 ms, for our protocols with TRs from 3 to 3.5 s, to provide high SNR-efficiency for our chosen ACE-EPTI protocols.

Figure 4 presents the SNR maps calculated from the anthropomorphic head phantom and in-vivo data acquired using ACE-EPTI and ss-EPI, at 1.5-mm isotropic (Fig. 4a), and 0.86 × 0.86 × 3 mm^3^ resolutions (Fig. 4b). The ss-EPI data were acquired with 3 averages to match its scan time with the 3-shot ACE-EPTI. As shown by the SNR maps, significantly higher SNR is provided by the echo-combined ACE-EPTI images when compared to the image generated by ss-EPI in both phantom and in-vivo acquisitions. For the 1.5-mm protocol, the mean SNR value (listed in each map) of ACE-EPTI is 57% higher than ss-EPI in phantom, and 36% higher in vivo. For the 0.86-mm protocol, the mean SNR of ACE-EPTI is 29% higher than ss-EPI in phantom, and 27% higher in vivo. The SNR gain is higher at 1.5-mm because of the much longer ETL of ACE-EPTI when compared to ss-EPI (56 ms vs. 20 ms), while the SNR gain for the high-resolution protocol is mainly from the shorter TE achieved through echo-train shifting. In addition to the echo-combined images, the presented example single-echo images from the ACE-EPTI acquisitions also show higher SNR than ss-EPI, validating its high efficiency for multi-echo imaging.

**Figure 4:**
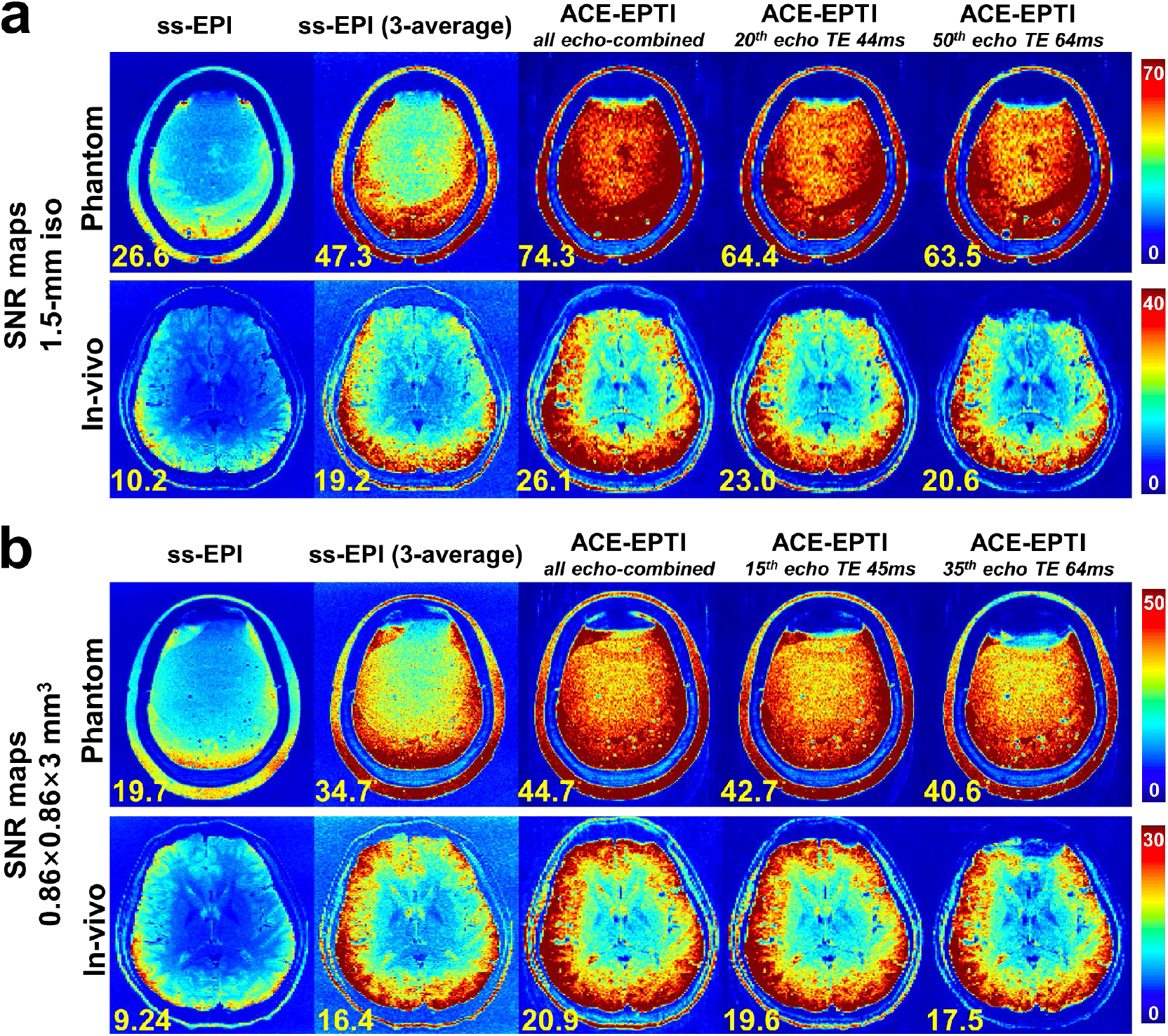
SNR maps of the ACE-EPTI and ss-EPI in both phantom and *in-vivo* experiments, at (a) 1.5-mm isotropic resolution, and (b) 0.86 × 0.86 × 3 mm^3^ resolution. SNR maps of 5 cases were shown, including (i) ss-EPI, (ii) ss-EPI with 3 averages, (iii) all-echo combined images of ACE-EPTI, (iv) single-echo image at TE = 44 or 45 ms of ACE-EPTI, and (v) single-echo image at TE = 64 ms of ACE-EPTI. The mean SNR values are listed in each map at the bottom left.

The effectiveness of the proposed phase correction framework is demonstrated through the reconstruction of an example in-vivo dMRI dataset shown in Fig. 5. Figure 5a compares the reconstructed non-DW image with and without the odd-even echo phase correction, as well as the estimated phase difference between the odd and even echoes. Based on the estimation, the major part of the odd-even echo phase difference is a 1D linear distribution along *x*, and the residual 2D phase difference after removing this 1D phase is on a much lower level (5-10 times smaller). After correcting for the phase difference, the ghosting-like artifacts around the edge (yellow arrows) is eliminated. The evaluation of the proposed self-navigated shot-to-shot phase correction is shown in Fig. 5b. Before correction, severe artifacts appear in all of the imaging slices due to the phase variation across the different shots (shown at the bottom). These artifacts are effectively removed after correction using the self-estimated phase, resulting in high quality diffusion images. Figure 5c shows the tracking of global *B*_0_ drifts obtained from the updated *B*_0_ maps across all the dynamics, where an increasing trend up to ~30 Hz is observed during the 20-minute scan. The estimated *B*_0_ change of the 40^th^ dynamic is presented in the figure as an example, and the corresponding reconstructed images with and without *B*_0_ updating are shown that demonstrates the importance of *B*_0_ updating for accurate image recovery (Fig. 5d).

**Figure 5:**
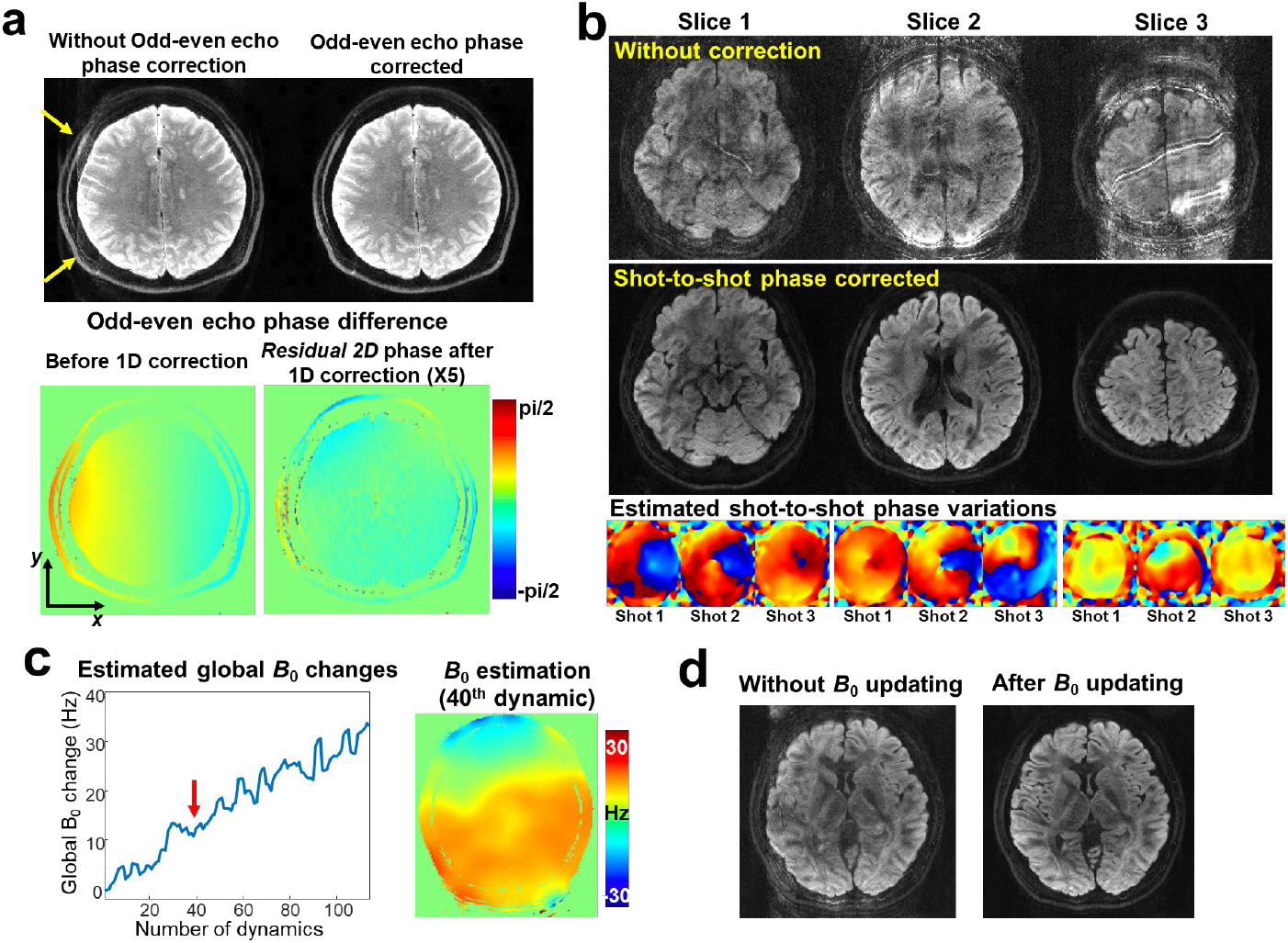
The evaluation of the proposed phase correction framework using *in-vivo* dMRI data. (a) Comparison of the reconstructed images with and without the odd-even echo phase correction, and the estimated phase difference between odd and even echoes before and after removing the 1D linear phase along *x*. Note that the residual 2D phase difference after removing 1D linear phase is magnified by 5 times. (b) The validation of the proposed self-navigated shot-to-shot phase correction. Reconstructed images before and after the shot-to-shot phase variation corrections are presented across 3 representative example slices, together with the estimated phase variation maps across the 3 shots. (c) The estimation of global *B*_0_ drifts obtained from the updated *B*_0_ maps across all of the dynamics during a 20-minute dMRI scan, and an example estimated *B*_0_ change map at the 40-th dynamic (including drifts and eddy currents). (d) The reconstructed images with and without the use of *B*_0_ updating at the 40-th dynamic.

The ability of ACE-EPTI to achieve distortion-free imaging is validated in Fig. 6, with the absent of susceptibility-induced (Fig. 6a) and eddy-currents distortions (Fig. 6b). Apparent distortions in the temporal and frontal lobes are observed in ss-EPI images (red arrows), even when an in-plane acceleration factor of 4 is used, and ghosting artifacts are also present indicated by the yellow arrows. The ACE-EPTI images are free from such distortion and artifacts, providing near identical image contours (red lines) to the TSE reference images. The dynamic image distortion due to eddy currents is also evaluated in Fig. 6b. The zoomed-in view of the ss-EPI images (with motion-correction but without eddy-current correction) across different diffusion directions shows clear up and down shifts of the edge. In contrast, ACE-EPTI provides stable and perfectly aligned images without dynamic shifts, demonstrating its robustness to eddy currents distortion.

**Figure 6:**
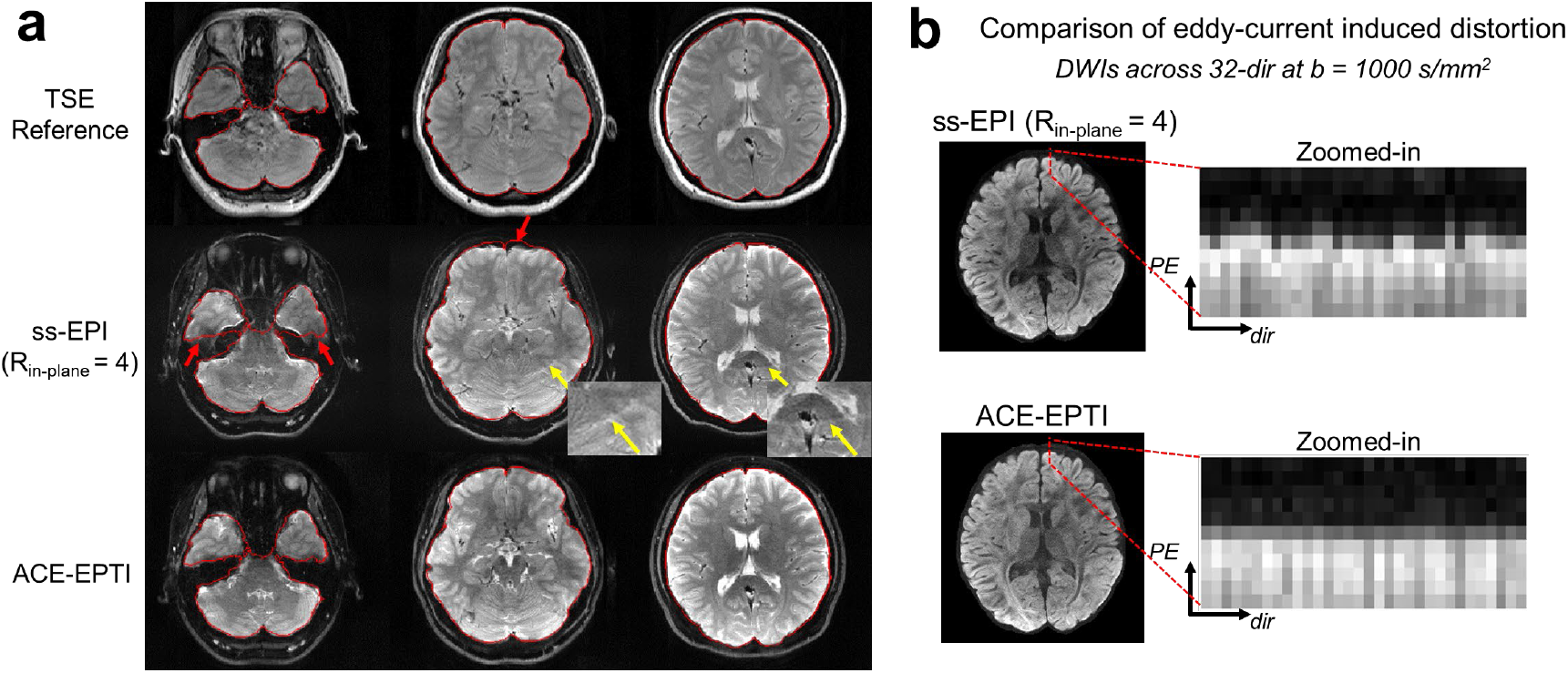
The evaluation of image distortion of ss-EPI and ACE-EPTI. (a) Comparison of susceptibility-induced distortions, where ACE-EPTI obtains near identical image contour to the TSE reference without distortion, while ss-EPI suffers from severe image distortions (red arrows), even with an in-plane acceleration factor of 4. Ghosting artifacts also appear in ss-EPI indicated by the zoomed-in areas (yellow arrows). (b) Comparison of dynamic distortions caused by eddy currents across diffusion directions. An anterior edge area of the brain (dashed red line) is zoomed-in along the PE direction with dynamics across all diffusion directions, showing up/down shifts of the ss-EPI images, and static and perfectly aligned edges of the ACE-EPTI, demonstrating the robustness of ACE-EPTI to eddy currents.

Figure 7 presents the diffusion images across different slices acquired by ACE-EPTI at 0.86-mm in-plane resolution. DWIs from 4 representative diffusion directions and 4 different TEs are shown, with high image quality and SNR. The calculated colored-FA maps obtained from the same data are shown in Fig. 8. Within the same acquisition time, FA maps from ACE-EPTI shows higher image quality with clearer and more consistent structures than ss-EPI. This is highlighted by the zoomed-in areas and yellow arrows. The improved quality of DTI can be attributed to the higher SNR and distortion and blurring-free feature of ACE-EPTI.

**Figure 7:**
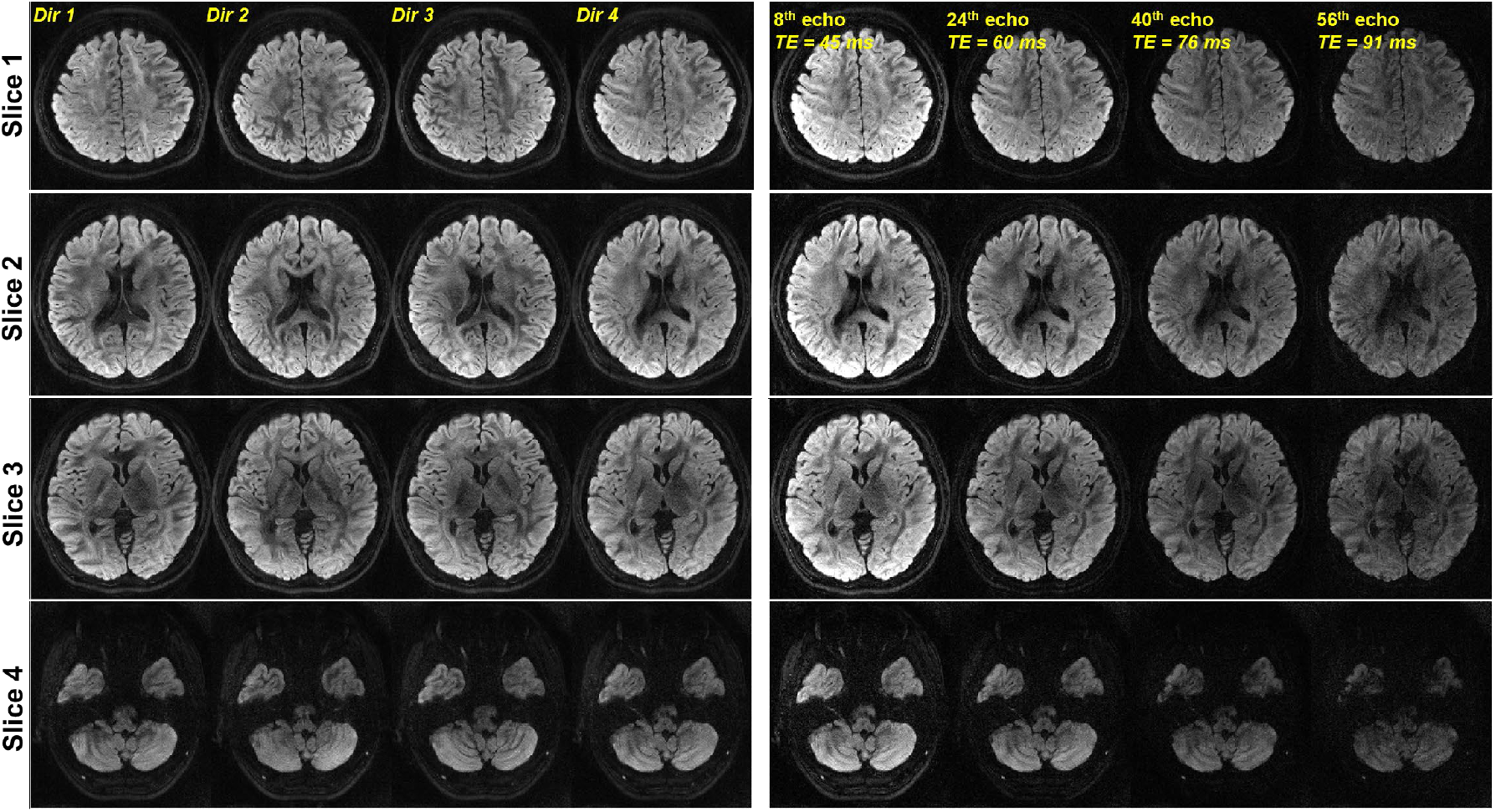
Representative diffusion weighted images from 4 diffusion directions and 4 different TEs acquired by ACE-EPTI at 0.86-mm in-plane resolution.

**Figure 8:**
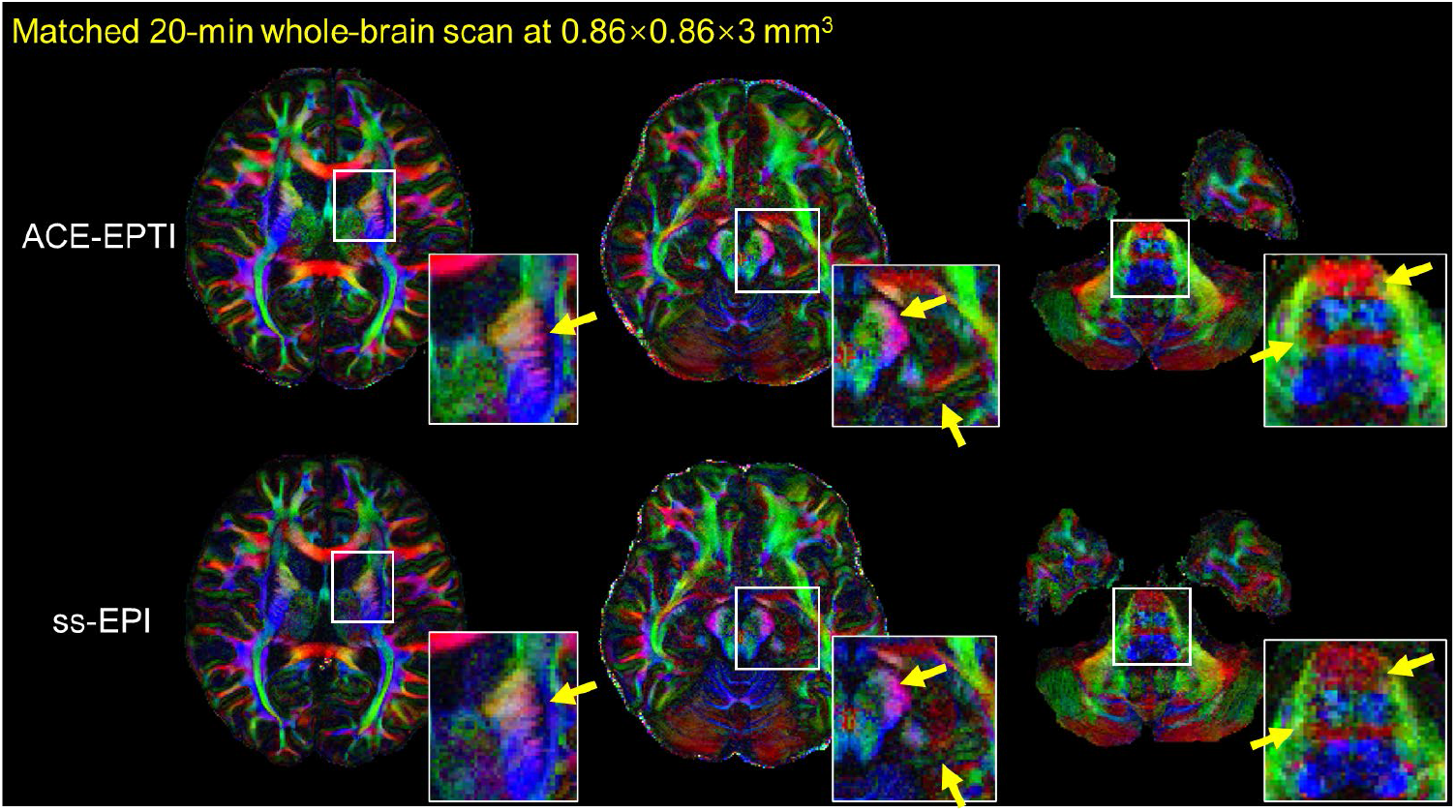
Comparison of colored-FA maps from ACE-EPTI and ss-EPI with matched scan time at 0.86 × 0.86 × 3 mm^3^. ACE-EPTI provides higher quality FA maps and more consistent structures than ss-EPI highlighted by the zoomed-in areas and yellow arrows.

The use of multi-echo images provided by ACE-EPTI is preliminarily evaluated in Fig. 9 at 0.86-mm in-plane resolution, and Fig. 10 at 1.5-mm isotropic resolution. Figure 9a shows the fitted T_2_* maps using averaged non-DW images and diffusion images, where a more homogeneous and lower value is observed from the DW-T_2_*, which could be caused by the significant reduction of the extracellular water signal after diffusion weighting, whose T_2_* should be longer than that of axonal and myelin water. Figure 9b shows example cFA maps at different TEs. Using the multi-TE data, different ODFs generated using data at different TEs are shown in Fig. 9c, where a fiber along the anterior-posterior direction (green-colored) which are present (white arrow) in the late TE is not visible in the early TE. Figure 10 shows the results of the 1.5-mm isotropic data with 3 different *b*-values (0, 1000, 2000 s/mm^2^). The fitted T_2_* maps using the mean multi-echo images (Fig. 10a) at three *b*-values are presented (Fig. 10b), and lower values can be seen under higher *b* values, which is consistent with the previous observation from Fig. 9a, potentially caused by the reduction of extracellular water signal. Figure 10c shows whole-brain cFA maps in 3 orthogonal views, and ODFs at different TEs of a zoomed-in area within the semiovale are presented in Fig. 10d. The ODFs show substantial differences across TEs, for example, a left-right fiber (red-colored) visible in early TEs gradually disappears in the later TEs (white arrows), indicating the additional information provided from data at multiple TEs.

**Figure 9:**
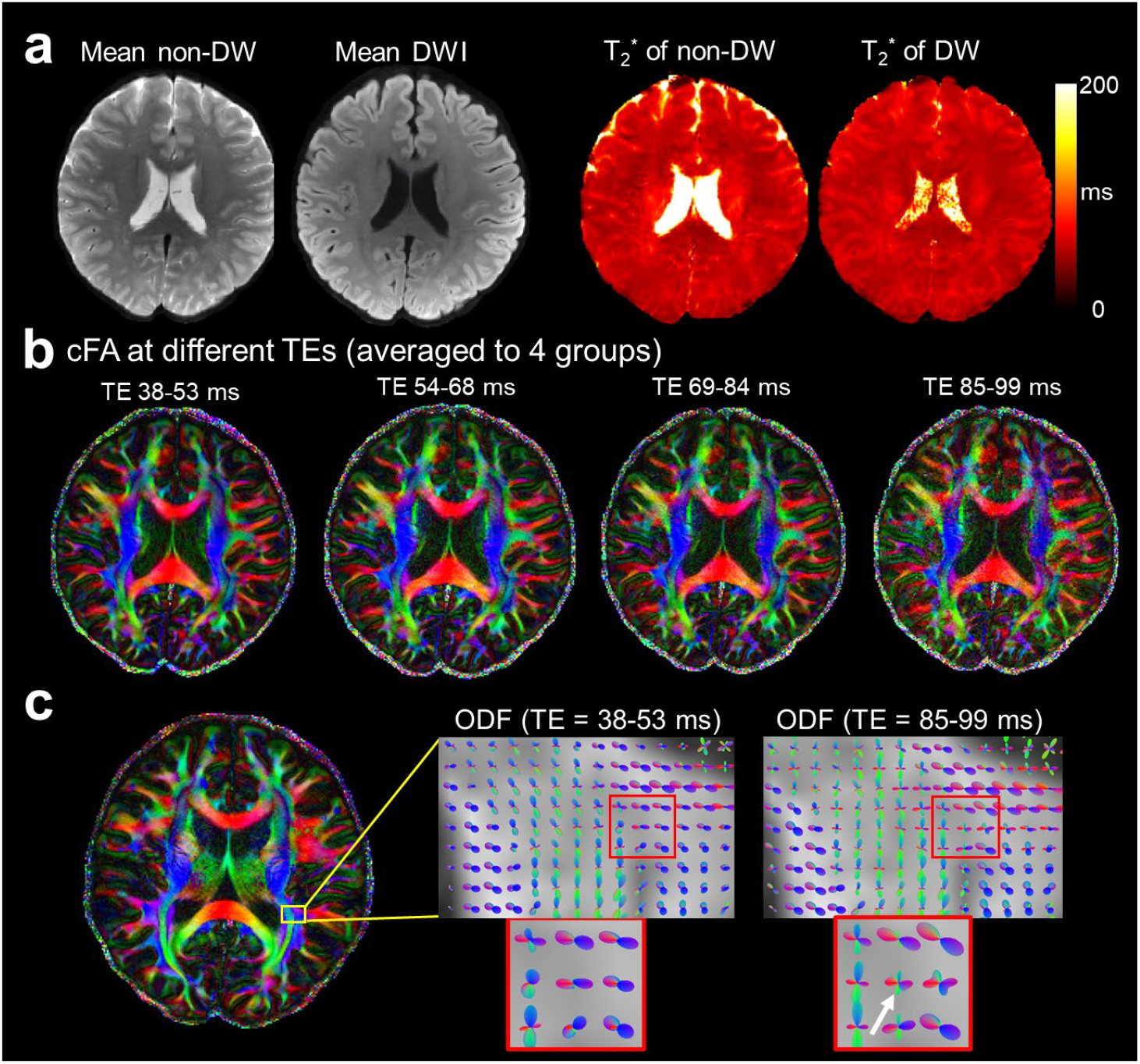
Results of the 0.86-mm in-plane experiment using ACE-EPTI. (a) Mean non-DW and DW images, and the fitted T_2_* maps from the mean images with and without diffusion weighting. (b) Colored-FA maps at different TEs. (c) Calculated ODFs at early (38-53 ms) and late TEs (85-99 ms). In the ODF from the late TEs, a connection along the anterior-posterior direction (green-colored) is present (white arrow) which is not visible in the early TE data.

**Figure 10:**
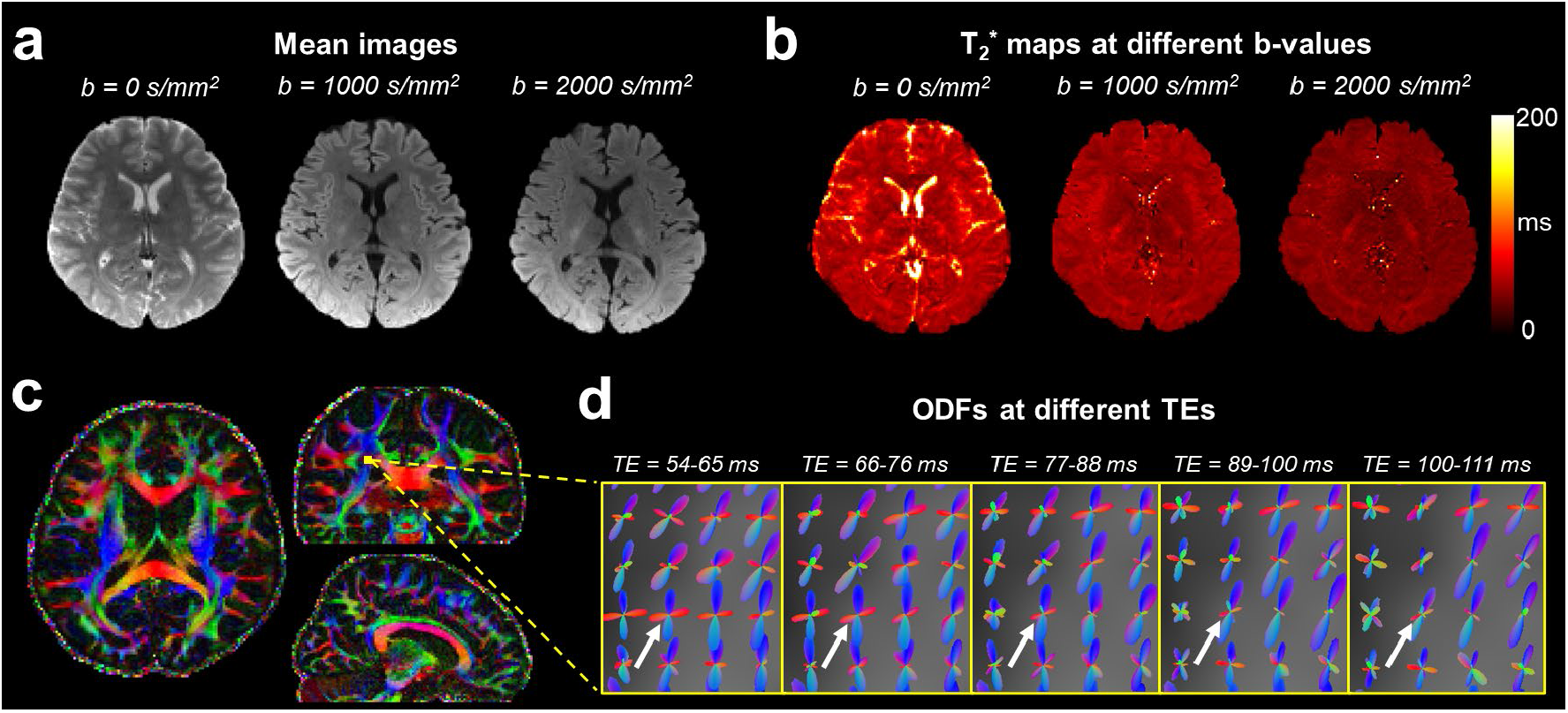
Results of the 1.5-mm isotropic experiment with 3 different b-values (0, 1000, 2000 s/mm^2^) acquired by ACE-EPTI. (a) Mean images at 3 different b-values. (b) T_2_* maps fitted from the mean multi-echo images at three b-values. (c) Three orthogonal views of the whole-brain cFA maps. (d) ODFs at different TEs of a zoomed-in area located at the semiovale. The white arrows highlight a left-right fiber (red-colored) which is visible in the early TEs but gradually disappears in the later TEs.

## Discussion

The results have demonstrated the improved SNR efficiency of ACE-EPTI over conventional ss-EPI, and also the effectiveness of the phase-corrected subspace reconstruction framework to provide artifact-free diffusion images. In addition, the fast 3-shot acquisition enabled by the variable density spatiotemporal encoding reduces motion sensitivity. Both susceptibility and eddy currents induced distortions are eliminated in the time-resolved multi-echo images, providing high anatomical integrity without the need for post-processing corrections.

The higher SNR efficiency of ACE-EPTI is attributed to the minimized TE and optimized ETL. At low resolutions, where the typical EPI readout window is short, the optimized ETL can improve the SNR efficiency more by using a longer readout to acquire more signals for noise averaging. Although the ETL of ss-EPI can be increased by reducing the in-plane acceleration and partial Fourier factors, the resultant longer TE and stronger blurring and distortion artifacts would compromise the SNR and image quality. At high resolution, the benefit of TE reduction using echo-train shifting over the conventional acquisition is more prominent, since the long echo train before the spin-echo with large echo spacing will lead to long TE in ss-EPI. The benefit of minimizing the TE has also been studied in many previous works, such as center-out spiral (49,50), and center-out EPI (51). One challenge of using a center-out trajectory in conventional single-shot acquisition is that the fast T_2_* decay would lead to more image blurring that needs to be corrected. This problem is fully addressed in ACE-EPTI which time resolves the signal evolution, providing images free from the signal-decay related blurring.

The distortion-free feature is another important advantage of ACE-EPTI. The time-resolved images are not only free from the susceptibility distortion, but also dynamic eddy currents across different diffusion directions as demonstrated in Fig. 6. There are still minor differences in the image contour of the ACE-EPTI and TSE images in Fig. 5a, which may be caused by small displacements between the two scans. In the time-resolving approach, the dynamic change of *B*_0_ from eddy currents across different diffusion directions would not lead to distortion, however, the reconstruction accuracy can be compromised. By updating the *B*_0_ change of every diffusion-encoded volume (Fig. 2), the *B*_0_ change due to eddy currents and drift can be estimated and corrected to ensure accurate reconstruction (Fig. 5 c&d). In addition to eddy currents, the odd-even echo phase difference and shot-to-shot phase variations are also effectively corrected by the proposed phase correction framework, eliminating potential image artifacts (Fig. 5 a&b).

The capability to resolve multi-echo images in ACE-EPTI provides an efficient acquisition for diffusion-relaxometry and TE-dependent diffusion analysis. Many previous studies have shown that the richer information provided by a joint diffusion-relaxometry acquisition could help disentangle more complicated tissue composition (8,10), detect unique features of diseases (9), and might be useful in tractography (52). The potential use of the time-resolved multi-echo images from ACE-EPTI is preliminary evaluated in this study, showing that multi-TE ODFs provide richer fiber orientation information. The T_2_* values are also different with different diffusion weightings, which could be caused by the change of signal contribution between axonal and extracellular water compartments under different diffusion weightings. To obtain the T1 relaxation information (11) together with T_2_/T_2_*, the efficient ACE-EPTI acquisition can be further combined with an inversion-recovery preparation. To provide shorter TRs and higher SNR-efficiency for whole-brain imaging, SMS acquisition has been integrated into the ACE-EPTI, and shown consistent high-quality results. To further improve the efficiency for a high isotropic resolution acquisition, simultaneous multi-slab acquisition such as gSlider (53–55) can also be combined with ACE-EPTI in the future. Additionally, ACE-EPTI can be also applied to other imaging applications outside of dMRI, such as multi-echo fMRI (56,57) and T_2_* mapping that has been previously studied using EPTI (28,58), with effective correction of shot-to-shot variation and a higher temporal resolution.

## Conclusions

ACE-EPTI was demonstrated to be a powerful and versatile technique for high-resolution diffusion MRI and diffusion-relaxometry. With a fast 3-shot acquisition scheme, ACE-EPTI is able to provide high SNR-efficiency, high image quality without distortion, blurring, and ghost artifacts, effective self-navigated phase correction, and capability for time-resolved multi-echo imaging.

## Acknowledgements

This work was supported by NIH NIBIB (R01-EB020613, R01-EB019437, R01-MH116173, P41-EB015896 and U01-EB025162) and by the MGH/HST Athinoula A. Martinos Center for Biomedical Imaging; and was made possible by the resources provided by NIH Shared Instrumentation Grants S10-RR023401, S10-RR023043, and S10-RR019307. We thank Dr. Lipeng Ning and Dr. Yogesh Rathi for the helpful discussion on diffusion-relaxometry.

## References

1. Le Bihan D, Breton EJCrdlAdsS, Mécanique, Physique, Chimie, Sciences de l’univers, Sciences de la Terre. Imagerie de diffusion in vivo par résonance magnétique nucléaire. 1985;301(15):1109–1112.

2. Setsompop K, Kimmlingen R, Eberlein E, Witzel T, Cohen-Adad J, McNab JA, Keil B, Tisdall MD, Hoecht P, Dietz P, Cauley SF, Tountcheva V, Matschl V, Lenz VH, Heberlein K, Potthast A, Thein H, Van Horn J, Toga A, Schmitt F, Lehne D, Rosen BR, Wedeen V, Wald LL. Pushing the limits of in vivo diffusion MRI for the Human Connectome Project. NeuroImage 2013;80:220–233.

3. McNab JA, Edlow BL, Witzel T, Huang SY, Bhat H, Heberlein K, Feiweier T, Liu K, Keil B, Cohen-Adad J, Tisdall MD, Folkerth RD, Kinney HC, Wald LL. The Human Connectome Project and beyond: initial applications of 300 mT/m gradients. NeuroImage 2013;80:234–245.

4. Miller KL, Alfaro-Almagro F, Bangerter NK, Thomas DL, Yacoub E, Xu J, Bartsch AJ, Jbabdi S, Sotiropoulos SN, Andersson JL, Griffanti L. Multimodal population brain imaging in the UK Biobank prospective epidemiological study. Nat Neurosci 2016;19(11): 1523–1536.

5. Holdsworth SJ, O’Halloran R, Setsompop K. The quest for high spatial resolution diffusion-weighted imaging of the human brain in vivo. NMR Biomed 2019;32(4):e4056.

6. McNab JA, Polimeni JR, Wang R, Augustinack JC, Fujimoto K, Stevens A, Janssens T, Farivar R, Folkerth RD, Vanduffel W. Surface based analysis of diffusion orientation for identifying architectonic domains in the in vivo human cortex. Neuroimage 2013;69:87–100.

7. Ji Y, Gagoski B, Hoge WS, Rathi Y, Ning L. Accelerated diffusion and relaxation-diffusion MRI using time-division multiplexing EPI. Magn Reson Med 2021.

8. Ning L, Gagoski B, Szczepankiewicz F, Westin CF, Rathi Y. Joint RElaxation-Diffusion Imaging Moments to Probe Neurite Microstructure. IEEE Trans Med Imaging 2020;39(3):668–677.

9. Slator PJ, Hutter J, Palombo M, Jackson LH, Ho A, Panagiotaki E, Chappell LC, Rutherford MA, Hajnal JV, Alexander DC. Combined diffusion-relaxometry MRI to identify dysfunction in the human placenta. Magn Reson Med 2019;82(1):95–106.

10. Veraart J, Novikov DS, Fieremans E. TE dependent Diffusion Imaging (TEdDI) distinguishes between compartmental T2 relaxation times. NeuroImage 2018;182:360–369.

11. Hutter J, Slator PJ, Christiaens D, Teixeira R, Roberts T, Jackson L, Price AN, Malik S, Hajnal JV. Integrated and efficient diffusion-relaxometry using ZEBRA. Sci Rep 2018;8(1): 15138.

12. Sodickson DK, Manning WJ. Simultaneous acquisition of spatial harmonics (SMASH): Fast imaging with radiofrequency coil arrays. Magn Reson Med 1997;38(4):591–603.

13. Pruessmann KP, Weiger M, Scheidegger MB, Boesiger P. SENSE: sensitivity encoding for fast MRI. Magn Reson Med 1999;42(5):952–962.

14. Griswold MA, Jakob PM, Heidemann RM, Nittka M, Jellus V, Wang J, Kiefer B, Haase A. Generalized autocalibrating partially parallel acquisitions (GRAPPA). Magn Reson Med 2002;47(6):1202–1210.

15. Bammer R, Keeling SL, Augustin M, Pruessmann KP, Wolf R, Stollberger R, Hartung HP, Fazekas F. Improved diffusion-weighted single-shot echo-planar imaging (EPI) in stroke using sensitivity encoding (SENSE). Magn Reson Med 2001;46(3):548–554.

16. Butts K, de Crespigny A, Pauly JM, Moseley M. Diffusion-weighted interleaved echo-planar imaging with a pair of orthogonal navigator echoes. Magn Reson Med 1996;35(5):763–770.

17. Chen NK, Guidon A, Chang HC, Song AW. A robust multi-shot scan strategy for high-resolution diffusion weighted MRI enabled by multiplexed sensitivity-encoding (MUSE). NeuroImage 2013;72:41–47.

18. Porter DA, Heidemann RM. High resolution diffusion-weighted imaging using readout-segmented echo-planar imaging, parallel imaging and a two-dimensional navigator-based reacquisition. Magn Reson Med 2009;62(2):468–475.

19. Holdsworth SJ, Skare S, Newbould RD, Bammer R. Robust GRAPPA-accelerated diffusion-weighted readout-segmented (RS)-EPI. Magn Reson Med 2009;62(6): 1629–1640.

20. Dong Z, Wang F, Ma X, Zhang Z, Dai E, Yuan C, Guo H. Interleaved EPI diffusion imaging using SPIRiT - based reconstruction with virtual coil compression. Magn Reson Med 2018;79(3):1525–1531.

21. Dong Z, Wang F, Ma X, Dai E, Zhang Z, Guo H. Motion-corrected k-space reconstruction for interleaved EPI diffusion imaging. Magn Reson Med 2018;79(4): 1992–2002.

22. Liu C, Bammer R, Kim Dh, Moseley ME. Self - navigated interleaved spiral (SNAILS): Application to high-resolution diffusion tensor imaging. Magn Reson Med 2004;52(6):1388–1396.

23. Robson MD, Gore JC, Constable RT. Measurement of the point spread function in MRI using constant time imaging. Magn Reson Med 1997;38(5):733–740.

24. Zeng H, Constable RT. Image distortion correction in EPI: comparison of field mapping with point spread function mapping. Magn Reson Med 2002;48(1): 137–146.

25. Zaitsev M, Hennig J, Speck O. Point spread function mapping with parallel imaging techniques and high acceleration factors: fast, robust, and flexible method for echo-planar imaging distortion correction. Magn Reson Med 2004;52(5): 1156–1166.

26. In MH, Posnansky O, Speck O. High-resolution distortion-free diffusion imaging using hybrid spin-warp and echo-planar PSF-encoding approach. NeuroImage 2017;148:20–30.

27. Dong Z, Wang F, Reese TG, Manhard MK, Bilgic B, Wald LL, Guo H, Setsompop K. Tilted-CAIPI for highly accelerated distortion-free EPI with point spread function (PSF) encoding. Magn Reson Med 2019;81(1):377–392.

28. Wang F, Dong Z, Reese TG, Bilgic B, Katherine Manhard M, Chen J, Polimeni JR, Wald LL, Setsompop K. Echo planar time-resolved imaging (EPTI). Magn Reson Med 2019;81(6):3599–3615.

29. Wang F, Dong Z, Reese TG, Rosen B, Wald LL, Setsompop K. 3D Echo Planar Time-resolved Imaging (3D-EPTI) for ultrafast multi-parametric quantitative MRI. bioRxiv 2021.

30. Dong Z, Wang F, Chan KS, Reese TG, Bilgic B, Marques JP, Setsompop K. Variable flip angle echo planar time-resolved imaging (vFA-EPTI) for fast high-resolution gradient echo myelin water imaging. NeuroImage 2021;232:117897.

31. Fair MJ, Wang F, Dong Z, Reese TG, Setsompop K. Propeller echo-planar time-resolved imaging with dynamic encoding (PEPTIDE). Magn Reson Med 2020;83(6):2124–2137.

32. Fair MJ, Liao C, Manhard MK, Setsompop K. Diffusion-PEPTIDE: Distortion- and blurring-free diffusion imaging with self-navigated motion-correction and relaxometry capabilities. Magn Reson Med 2021; 85(5):2417–2433.

33. Eichner C, Paquette M, Mildner T, Schlumm T, Pleh K, Samuni L, Crockford C, Wittig RM, Jager C, Moller HE, Friederici AD, Anwander A. Increased sensitivity and signal-to-noise ratio in diffusion-weighted MRI using multi-echo acquisitions. NeuroImage 2020;221:117172.

34. Setsompop K, Gagoski BA, Polimeni JR, Witzel T, Wedeen VJ, Wald LL. Blipped-controlled aliasing in parallel imaging for simultaneous multislice echo planar imaging with reduced g-factor penalty. Magn Reson Med 2012;67(5):1210–1224.

35. Liang Z-P. Spatiotemporal imagingwith partially separable functions. 2007. IEEE. p 988–991.

36. Tamir JI, Uecker M, Chen W, Lai P, Alley MT, Vasanawala SS, Lustig M. T2 shuffling: Sharp, multicontrast, volumetric fast spin-echo imaging. Magn Reson Med 2017;77(1): 180–195.

37. Dong Z, Wang F, Reese TG, Bilgic B, Setsompop K. Echo planar time-resolved imaging with subspace reconstruction and optimized spatiotemporal encoding. Magn Reson Med 2020;84(5):2442–2455.

38. Lam F, Liang ZP. A subspace approach to high-resolution spectroscopic imaging. Magn Reson Med 2014;71(4):1349–1357.

39. Zhang T, Pauly JM, Levesque IR. Accelerating parameter mapping with a locally low rank constraint. Magn Reson Med 2015;73(2):655–661.

40. Tamir JI, Ong F, Cheng JY, Uecker M, Lustig M. Generalized magnetic resonance image reconstruction using the Berkeley advanced reconstruction toolbox. ISMRM Workshop on Data Sampling & Image Reconstruction 2016.

41. Uecker M, Ong F, Tamir JI, Bahri D, Virtue P, Cheng JY, Zhang T, Lustig M. Berkeley advanced reconstruction toolbox. . In Proc Intl Soc Mag Reson Med 2015;p 2486.

42. Jenkinson M, Bannister P, Brady M, Smith S. Improved optimization for the robust and accurate linear registration and motion correction of brain images. Neuroimage 2002;17(2):825–841.

43. Jenkinson M, Smith S. A global optimisation method for robust affine registration of brain images. Med Image Anal 2001;5(2):143–156.

44. Jenkinson M, Beckmann CF, Behrens TE, Woolrich MW, Smith SM. Fsl. NeuroImage 2012;62(2):782–790.

45. Smith SM, Jenkinson M, Woolrich MW, Beckmann CF, Behrens TE, Johansen-Berg H, Bannister PR, De Luca M, Drobnjak I, Flitney DE. Advances in functional and structural MR image analysis and implementation as FSL. Neuroimage 2004;23:S208–S219.

46. Smith SM. Fast robust automated brain extraction. Hum Brain Mapp 2002;17(3):143–155.

47. Tournier JD, Calamante F, Connelly A. MRtrix: Diffusion tractography in crossing fiber regions. International Journal of Imaging Systems and Technology 2012;22(1):53–66.

48. Andersson JL, Sotiropoulos SN. An integrated approach to correction for off-resonance effects and subject movement in diffusion MR imaging. NeuroImage 2016;125:1063–1078.

49. Lee Y, Wilm BJ, Brunner DO, Gross S, Schmid T, Nagy Z, Pruessmann KP. On the signal-to-noise ratio benefit of spiral acquisition in diffusion MRI. Magn Reson Med 2021;85(4): 1924–1937.

50. Wilm BJ, Hennel F, Roesler MB, Weiger M, Pruessmann KP. Minimizing the echo time in diffusion imaging using spiral readouts and a head gradient system. Magn Reson Med 2020;84(6):3117–3127.

51. Chen X, Zhu A, Du YP. Center-out EPI (COEPI): A fast single-shot imaging technique with a short TE. Magn Reson Med 2020;84(2):787–799.

52. de Almeida Martins JP, Tax CMW, Reymbaut A, Szczepankiewicz F, Chamberland M, Jones DK, Topgaard D. Computing and visualising intra-voxel orientation-specific relaxation-diffusion features in the human brain. Hum Brain Mapp 2021;42(2):310–328.

53. Setsompop K, Fan Q, Stockmann J, Bilgic B, Huang S, Cauley SF, Nummenmaa A, Wang F, Rathi Y, Witzel T, Wald LL. High-resolution in vivo diffusion imaging of the human brain with generalized slice dithered enhanced resolution: Simultaneous multislice (gSlider-SMS). Magn Reson Med 2018;79(1):141–151.

54. Wang F, Bilgic B, Dong Z, Manhard MK, Ohringer N, Zhao B, Haskell M, Cauley SF, Fan Q, Witzel T, Adalsteinsson E, Wald LL, Setsompop K. Motion-robust sub-millimeter isotropic diffusion imaging through motion corrected generalized slice dithered enhanced resolution (MC-gSlider) acquisition. Magn Reson Med 2018;80(5):1891–1906.

55. Liao C, Bilgic B, Tian Q, Stockmann JP, Cao X, Fan Q, Iyer SS, Wang F, Ngamsombat C, Lo WC, Manhard MK, Huang SY, Wald LL, Setsompop K. Distortion-free, high-isotropic-resolution diffusion MRI with gSlider BUDA-EPI and multicoil dynamic B0 shimming. Magn Reson Med 2021;86(2):791–803.

56. Kundu P, Voon V, Balchandani P, Lombardo MV, Poser BA, Bandettini PA. Multi-echo fMRI: a review of applications in fMRI denoising and analysis of BOLD signals. Neuroimage 2017;154:59–80.

57. Kundu P, Inati SJ, Evans JW, Luh WM, Bandettini PA. Differentiating BOLD and non-BOLD signals in fMRI time series using multi-echo EPI. NeuroImage 2012;60(3):1759–1770.

58. Wang F, Dong Z, Wald LL, Polimeni JR, Setsompop K. Simultaneous pure T2 and varying T2’-weighted BOLD fMRI using Echo Planar Time-resolved Imaging (EPTI) for mapping laminar fMRI responses. bioRxiv 2021.

